# Cell wall composition in relation to photosynthesis across land plants’ phylogeny: crops as outliers

**DOI:** 10.1101/2024.08.07.606640

**Authors:** Margalida Roig-Oliver, Jaume Flexas, María José Clemente-Moreno, Marc Carriquí

## Abstract

In the present study, we combine published and novel data on cell wall composition and photosynthesis limitations, including data for all the major land plant’s phylogenetic groups. We provide novel evidence on the importance of cell wall composition in determining mesophyll conductance to CO_2_ diffusion (*g*_m_) across land plants’ phylogeny. We address the hypothesis that the pectin fraction of total major cell wall compounds is positively related to *g*_m_ and, consequently, to photosynthesis, when pooling species from across the entire phylogeny.

The role of cell wall composition in photosynthesis has only recently been proposed. Apparently contradictory results have been reported, but previous studies were often limited to single or closely related species. This is the very first report to show general relationships by considering species spanning the entire phylogeny of land plants.

This study identifies a clear biochemical basis—one that can be traced back to specific genes— for a large component of mesophyll conductance and, thus, photosynthetic capacity. It opens new avenues for improving the photosynthesis of terrestrial plants. Additionally, it suggests that current crops are already optimized and even uncoupled from these general relationships, raising questions about the regulation of *g*_m_ in crop species.

## The role of leaf functional traits determining photosynthesis

The *leaf economics spectrum* (LES) represents a complex framework of interconnected leaf traits related to carbon fixation and nutrient use across plant lineages (Wright *et al*., 2004), with net CO_2_ assimilation (*A*_N_) and leaf mass per area (LMA) as key traits (Onoda *et al*., 2017). Generally, LMA varies among plant species and negatively correlates with physiological traits such as *A*_N_ at a phylogenetic scale (Onoda *et al*., 2017). Similarly, lower LMA is linked with enhanced mesophyll conductance to CO_2_ diffusion (*g*_m_) (Onoda *et al*., 2017). This parameter, encompassing the CO_2_ pathway from substomatal cavities to Rubisco carboxylation sites in chloroplasts stroma, is recognized as a key trait regulating photosynthesis (Flexas *et al*., 2012; Flexas *et al*., 2018). However, the mechanistic basis by which *g*_m_ is regulated remains largely misunderstood (Evans *et al*., 2009; Flexas *et al*., 2012; Flexas *et al*., 2018; Mizokami *et al*., 2022).

In addition to LMA, anatomical properties such as the chloroplast surface area exposed to intercellular air spaces per unit leaf area (*S*_c/_*S*) and cell wall thickness (*T*_cw_)—often assumed to be positively linked with increased LMA—strongly influence *g*_m_ across different plant lineages (Carriquí *et al*., 2019; Flexas & Carriquí, 2020; Gago *et al*., 2019; Peguero-Pina *et al*., 2017; Tosens *et al*., 2016; Veromann-Jurgenson *et al*., 2020; Veromann-Jurgenson *et al*., 2017). Thick cell walls limit *g*_m_, suggesting a phylogenetic trend towards increasing *g*_m_ by reducing *T*_cw_ from mosses to angiosperms (Flexas & Carriquí, 2020; Gago *et al*., 2019). However, while the cell wall conductance to CO_2_ (*g*_cw_) strongly influences *g*_m_ and, consequently, photosynthesis, *T*_cw_— representing the apparent CO_2_ diffusion path length across the cell wall (Δ*L*_cw_)—is just one component. The other components are diffusivity (*D*), which is a function of leaf temperature, cell wall porosity (*p*_cw_), and cell wall tortuosity (τ_cw_) (Evans *et al*., 2009; Flexas *et al*., 2021; Terashima *et al*., 2011):

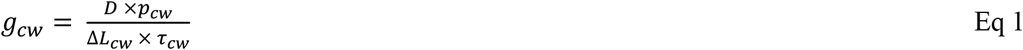

Although leaf temperature is unrelated to cell wall, porosity and tortuosity are theoretically influenced by cell wall composition, and even *T*_cw_ itself could be indirectly linked to it.

Recently, cell wall compositional traits were shown to be involved in *g*_m_ (Carriquí *et al*., 2020; Clemente-Moreno *et al*., 2019; Ellsworth *et al*., 2018; Roig-Oliver *et al*., 2020a,b, 2021a,b,c; Salesse-Smith *et al*., 2024). Although specific land plants lineages possess specific cell wall compositional characteristics (Popper *et al*., 2011; Sarkar *et al*., 2009; Sørensen *et al*., 2010), the cell wall is a three-dimensional framework mainly composed by cellulose microfibrils cross-linked to non-cellulosic polysaccharides (hemicelluloses), all embedded within a pectin matrix (Anderson & Kieber, 2020; Cosgrove, 2005; Somerville *et al*., 2004). This pectin network is believed to be a key structure determining several cell wall properties that could potentially affect CO_2_ diffusion, such as porosity, thickness and elasticity (Carriquí *et al*., 2020; Cosgrove, 2005; Flexas *et al*., 2021; Novakovic *et al*., 2018; Ochoa-Villareal *et al*., 2012; Roig-Oliver *et al*., 2020a,b; 2021b; 2022; Schiraldi *et al*., 2012; Weraduwage *et al*., 2016). In fact, recent studies provided empirical relationships between variations in pectin concentration and adjustments in *T*_cw_ and in the bulk modulus of elasticity (ε). However, these studies were performed either testing one or few species (Clemente-Moreno *et al*., 2019; Roig-Oliver *et al*., 2020a,b; 2021a,c) or varieties (Ellsworth *et al*., 2018; Roig-Oliver *et al*., 2021d, 2022) of angiosperms responding to environmental stresses or having specific genetic modifications, or being comparisons of non-stressed species within the same phylogenetic group (Carriquí *et al*., 2020; Roig-Oliver *et al*., 2021b). These studies suggested that either pectin concentration or the proportion between different cell wall components (i.e., the pectin to hemicelluloses and cellulose ratio) are crucial regulators of *g*_m_ and other leaf physiological traits, but in a variable manner (Carriquí *et al*., 2020; Roig-Oliver *et al*., 2020a; 2022).

To our knowledge, no previous study has addressed the implication of cell wall composition influencing both *g*_m_ and *T*_cw_ along land plants’ lineages. Thus, our aims were (1) to perform a meta-analysis using both literature and newly measured species in which cell wall composition, photosynthetic and leaf anatomical properties across species spanning from mosses to angiosperms are considered; and (2) to explore how cell wall compositional traits influence leaf anatomy and photosynthesis, being *T*_cw_, cell wall porosity and *g*_m_ key traits. Our main hypothesis is that cell wall composition and, particularly, the fraction of pectins over the other two major components (cellulose + hemicellulose) —as a proxy of cell wall porosity (and, perhaps, tortuosity) —contributes to determining *g*_m_ variations along land plants’ lineages.

## Variation along land plants’ phylogeny

Despite the modest number of species considered in our dataset, it still covers 70% or more of the published ranges (excluding the first and last decile) for each trait and phylogenetic group in 75% of the cases (Fig. S1). This comprehensive coverage ensures that our findings are representative and robust.

The exploration of the variation in structural and physiological traits, as well as those related to cell wall composition, across several land plant groups reveals significant insights (Fig. 1). Contrary to common assumptions, our findings suggest that there is no dependence of *T*_cw_ on LMA across phylogenetic scales (Fig. 1a-b). Instead, the patterns for *A*_N_ and *g*_m_ show remarkable similarities across different plant lineages, as already highlighted by Gago *et al*. (2019) and Huang *et al*. (2022) (Fig. 1c-d). This observation underscores the notion that limitations imposed by *g*_m_ are dominant throughout land plants’ phylogeny (Gago *et al*., 2019).

**Fig. 1.**
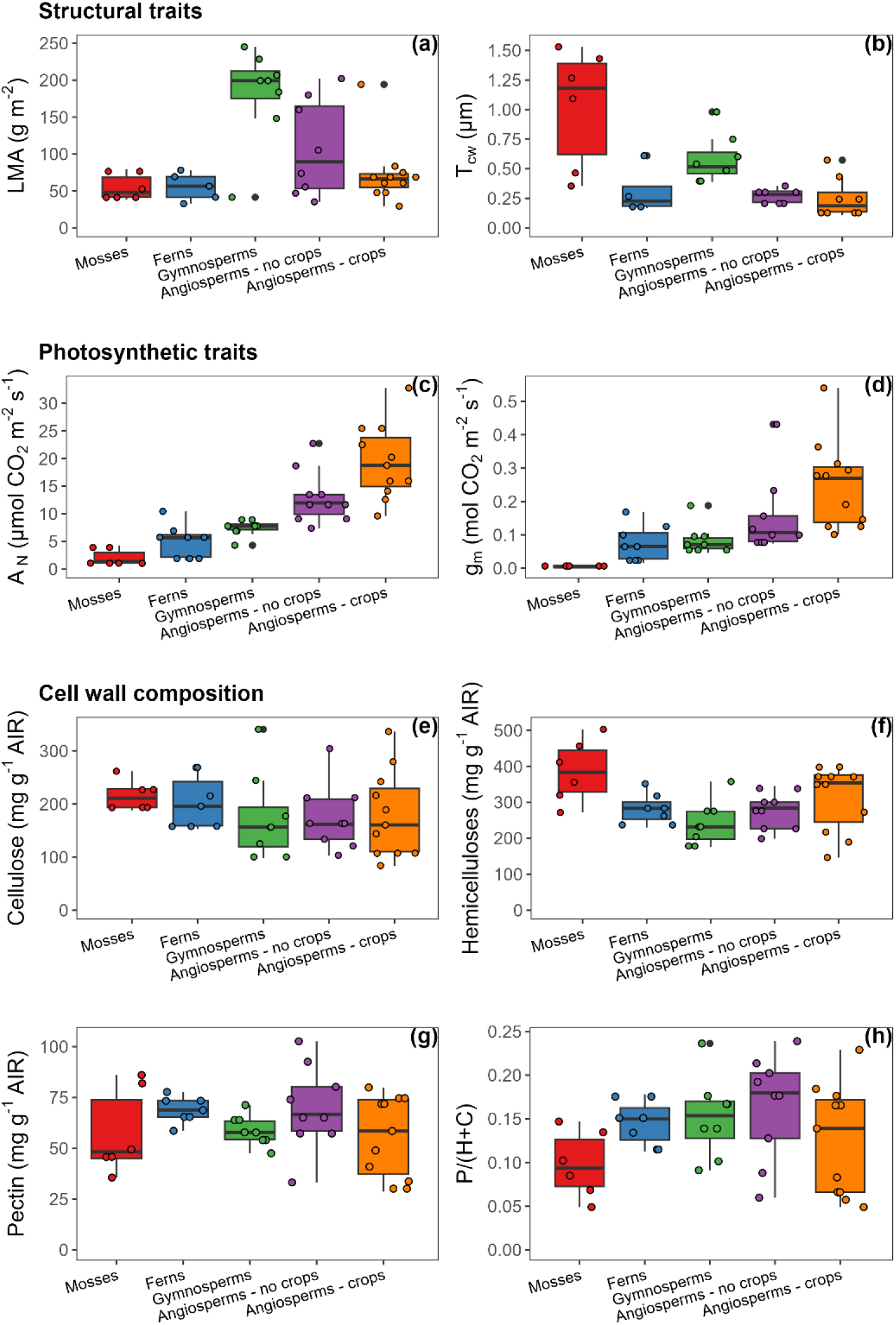
Phylogenetic trends in structural and physiological traits and in cell wall composition. Dot boxplots showing (a) leaf dry mass per unit area (LMA), (b) cell wall thickness (T_cw_), (c) net CO_2_ assimilation (A_N_), (d) mesophyll conductance (g_m_), (e) cellulose, (f), (g) pectin, and (h) pectin to cellulose and hemicellulose ratio (P/(C+H)) for mosses (n = 6), ferns (n = 8), gymnosperms (n = 8), wild angiosperms (n = 10), and angiosperm crops (n = 11). Data was compiled from Carriquí et al. (2020), Clemente-Moreno et al. (2019), Nadal et al. (2020), Nadal et al. (2023), Roig-Oliver et al. (2020b,c; 2021b,c; 2022), and from newly measured species.

The absence of similar trends in *T*_cw_ and LMA across groups suggests that these traits evolved independently, challenging traditional views on their interdependence. This finding emphasizes the complexity of plant adaptations, revealing that factors influencing *T*_cw_ do not necessarily correlate with leaf mass. Instead, the inverse but strong phylogenetic patterns observed in *T*_cw_ compared to *A*_N_ and *g*_m_ indicate that these structural and physiological traits are more closely linked, highlighting their functional relationship. Moreover, our analysis shows that there are no major differences among groups in main cell wall compounds contents or in the pectin to hemicellulose and cellulose ratio: intra-group variations appear to be more pronounced than inter-group variations (Fig. 1e-h).

## Dependency of photosynthesis on cell wall composition

Contrary to expectations based on Fig. 1, the analysis of the relationships between cell wall composition and *g*_m_ shown in Fig. 2 reveals several important insights, particularly when distinguishing between crop and non-crop species. As already shown (Evans, 2021; Flexas *et al*., 2021), *g*_m_ correlates negatively with *T*_cw_, which is a proxy for the effective path length across the cell wall, following an exponential decay function (Fig. 2a). In contrast, no significant correlation was found between *g*_m_ and LMA or leaf thickness, and the significant correlation with leaf density (LD) was driven by unistratose mosses only, which possess a very large density and low *g*_m_ (Fig. S2). Additionally, a significant positive linear correlation emerges between *g*_m_ and the pectin to cellulose plus hemicelluloses fraction (P/(C+H)), a proposed proxy for porosity and tortuosity (Carriquí *et al*., 2020; Flexas *et al*., 2021), thus confirming our hypothesis (Fig. 2b). However, this correlation is significant only when excluding crops (r^2^ = 0.35; *p* < 0.001), and it improves further when removing non-crop angiosperms (r^2^ = 0.57; *p* < 0.0001). Notably, a much stronger positive correlation between *g*_m_ and the ratio between P/(C+H) and *T*_cw_ was found (Fig. 2c), indicating the combined importance of path length—*T*_cw_—and porosity/tortuosity—P/(C+H). Again, this relationship is especially pronounced when crop species are excluded from our dataset.

**Fig. 2.**
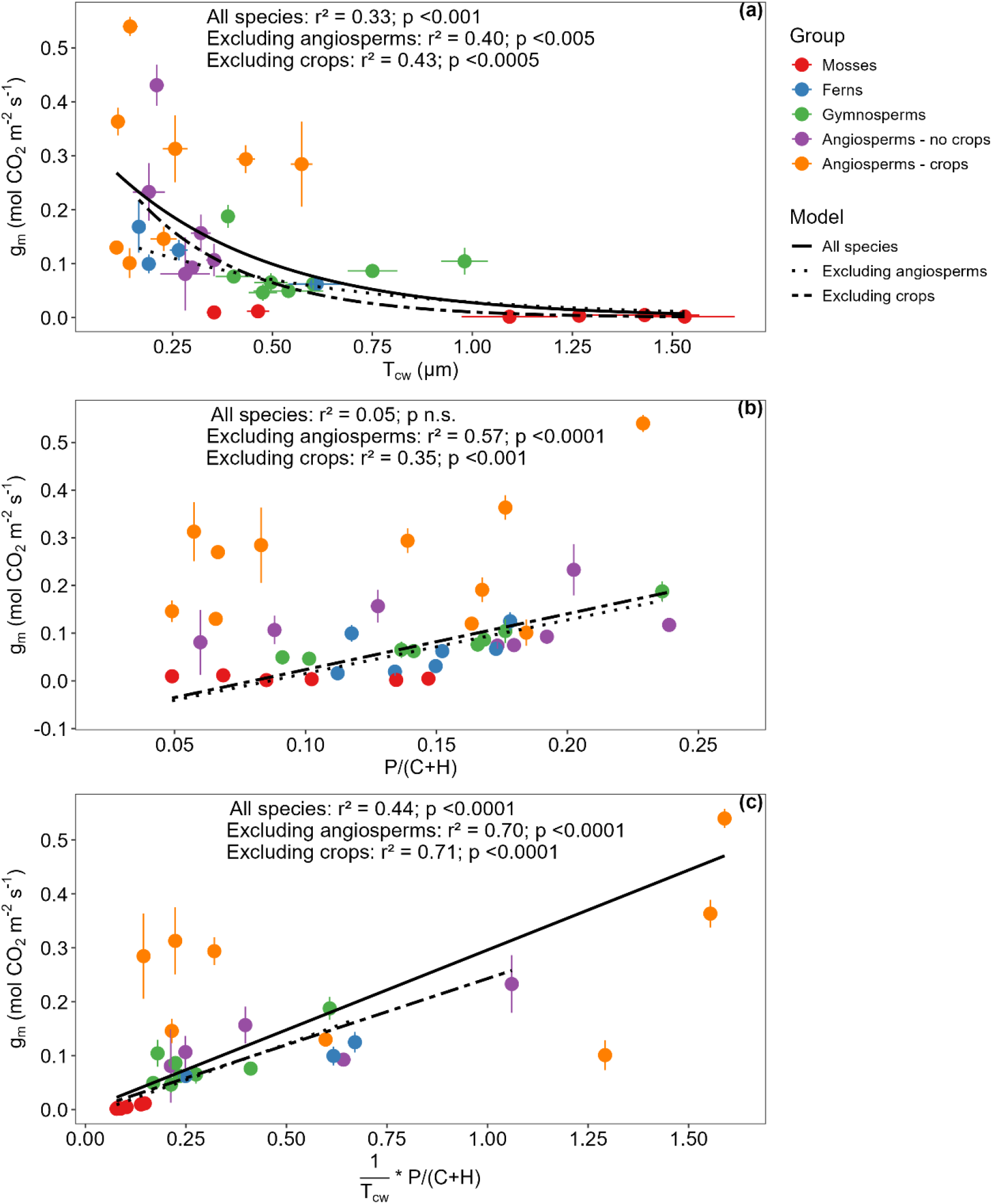
Relationships between mesophyll conductance (g_m_) and (a) cell wall thickness (T_cw_), (b) pectin to hemicellulose and cellulose ratio (P/(C+H)), and (c) 1/T_cw_ *P/(C+H). Lines represent exponential decay o linear fittings on data pooled across all phylogenetic groups, excluding crops and excluding all angiosperms. Significant lines in (a) are exponential decay regression lines and in (b, c) standardized major axis (SMA) regression lines. Data was compiled from Carriquí et al. (2020), Clemente-Moreno et al. (2019), Nadal et al. (2020), Nadal et al. (2023), Roig-Oliver et al. (2020b,c;2021b,c;2022), and from newly-measured species.

In non-crop species, the pectin fraction in relation to cellulose and hemicellulose plays a crucial role in determining how easily CO_2_ can diffuse through the leaf mesophyll. Conversely, the inclusion of crop species introduces a different dynamic. Crops have indirectly been selectively bred for enhanced photosynthetic efficiency and other agronomically valuable traits (Nadal & Flexas, 2019), which is the consequence of these plants having been derived from wild species with naturally high photosynthetic capacities (Gomez-Fernandez *et al*., 2024). They often possess very thin cell walls, which might have weakened the correlation between P/(C+H) and *g*_m_ because they are optimized for rapid growth and high yield rather than for natural selection pressures influencing photosynthetic efficiency (Xiong, 2023). Consequently, as shown in Fig. 2b, crops can achieve a large *g*_m_ despite not necessarily showing a large P/(C+H) ratio. This does not mean that cell wall thickness and/or composition does not affect *g*_m_ in crops. Indeed, improved photosynthesis and yield has been recently achieved by genetically manipulating cell walls (Salesse-Smith *et al*., 2024), and empirical relationships between *g*_m_ and cell wall composition have been shown in different studies subjecting crops to stress conditions (Roig-Oliver *et al*., 2020a,b; 2021a; 2022). However, it implies that the role in crops is lesser than in wild species and/or that crops may have compensating mechanisms for facilitating CO_2_ diffusion. The impact of crop selection on cell wall composition underscores the necessity of focusing on non-crop species to better understand the fundamental connections between cell wall components and photosynthetic traits.

From Eq. 1 the CO_2_ conductance across cell walls should depend on Δ*L*_cw_ and *p*_cw_ /τ_cw_ as the mixed variable, here approached as 1/*T*_cw_ * P/C+H). In fact, the correlation of this parameter with *g*_m_ across different plant groups becomes notably more significant, with all species having an r^2^ = 0.44, excluding angiosperms r^2^ = 0.70, and excluding crops r^2^ = 0.71, all with p < 0.0001. These results suggest that this combined variable significantly explains the variations in *g*_m_ (Fig. 2c), and emphasizes the relevant role that the pectin fraction plays in regulating *g*_m_ and, consequently, *A*_N_, in wild, non-domesticated plants. To fully comprehend this dependency, it is essential to explore how pectins influence various aspects of cell wall properties—such as porosity and thickness—that affect gas exchange. While the actual physical porosity should not be a major limitation given that CO_2_ molecules are small compared to cell wall pore sizes (Carpita *et al*., 1979; Flexas *et al*., 2021), pectins possess hydrocolloid characteristics and can retain several times their volume in water. Since CO_2_ diffuses in solution, the hydrophilic fraction of the pores could determine its diffusion (Flexas et al., 2021). Moreover, pectins can be under different methyl-esterification states and may interact with other chemical cell wall compounds, altogether affecting the ‘effective’ porosity and, perhaps, tortuosity (Flexas *et al*., 2021). By focusing on these fundamental connections, we can better direct efforts to improve photosynthesis in species with relatively thick cell walls and enhance their physiological performance under contrasting environmental conditions.

## Acknowledgments

This work was supported by projects PGC2018-093824-B-C41, PID2022-138424NB-I00 and PID2022-139455NB-C31 from the Ministerio de Economía y Competitividad (MINECO, Spain) and the ERDF (FEDER). MR-O was supported by a predoctoral fellowship (FPU16/01544) from Ministerio de Economía y Competitividad (MINECO, Spain). MC was supported by a Vicenç Mut 2022 postdoctoral fellowship (PD-047-2022) funded by Conselleria de Fons Europeus, Universitat i Cultura from Govern de les Illes Balears. MJC was supported by RYC2020-029602-I by MCIN/AEI/ 10.13039/501100011033 and by “ESF Investing in your future”. We are grateful to Drs. Cyril Douthe and Miquel Nadal for their help during gas exchange measurements, to Dr. Mateu Fullana-Pericàs and all the staff at the Serveis Cientificotècnics of the Universitat de les Illes Balears for their support and maintenance of gas-exchange/fluorescence equipment, to María Teresa Mínguez (Universitat de València – Secció Microscòpia Electrònica, SCSIE) and Dr Ferran Hierro (Universitat de les Illes Balears – Serveis Cientificotècnics) for technical support during microscopic analysis. Finally, we wish to thank Mr. Miquel Truyols and other collaborators of the UIB Experimental Field and Glasshouses, supported by the UIB Grant 15/2015.

## Competing interests

None declared.

## Author contributions

MC, MR-O and JF designed the research. MR-O measured the new studies presented in this study. MC performed the data analysis. MC, MR-O and JF wrote the manuscript. All authors corrected the manuscript.

## Data availability

All data is available in the Supplementary Materials.

## Supporting information

Additional Supporting Information may be found online in the Supporting Information section at the end of the article.

**Methods S1**. Methods used in this paper.

**Table S1**. Data used in this paper.

**Figure S1**. Mesophyll conductance in relation to leaf mass per area, leaf thickness and leaf density.

**Figure S2**. Dot boxplots showing dry mass-based cellulose fraction of alcohol-insoluble residues for the different land plant groups.

## SUPPORTING INFORMATION

**Methods S1** | Methods used in this paper

### Data compilation

We conducted a data compilation covering gas exchange, leaf anatomy and cell wall composition across a variety of land plant species. Our focus was on studies that met two key criteria: (1) they measured all relevant traits in the same plants and (2) they employed a consistent protocol for analyzing cell wall composition, ensuring the obtention of alcohol insoluble residues (AIR) within the range of previously reported values (Fig. S3). To enrich our dataset, we also conducted supplementary measurements on three wild angiosperm species and three ferns. The final dataset encompasses data from 21 angiosperms (including 10 wild species and 11 crops), 8 gymnosperms, 8 ferns and 6 mosses, all grown under non-stressful conditions (Table S1).

### Growing conditions of newly measured species

The angiosperm *Deschampsia antarctica* was sampled in the field on Livingston Island, South Shetland Islands, Antarctica (62º39’94’’S, 60º23’20’’W, 12 meters above sea level) during February 2020, which corresponds to the Antarctic summer. The other 5 species (the angiosperms *Arabidopsis thaliana* and *Camellia japonica*, and the ferns *Adiantum pedatum, Asplenium scolopendrium* and *Dryopteris erythrosora*) were cultivated at the Universitat de les Illes Balears in Mallorca, Spain. These plants were potted in a substrate mixture of peat and perlite in a 3:1 ratio. Apart from *Arabidopsis thaliana*, which was grown in a controlled chamber at 22 ºC with a photosynthetic photon flux density (PPFD) of 200-250 μmol m^-2^s^-1^ for 12 h of light followed by 12 h of darkness, all other plants were placed outdoors during the spring season to ensure optimal, non-stressing conditions. Plants were watered three times per week to maintain adequate moisture levels along the growing period.

### Photosynthetic characterization

Simultaneous gas exchange and fluorescence measurements were performed using an open gas exchange system with a 2 cm^2^ fluorescence chamber (Li-6400; Li-Cor Inc., Lincoln, USA). Measurements were taken on fully expanded leaves from five to six replicates per species. After reaching steady-state conditions, net CO_2_ assimilation (*A*_N_), stomatal conductance (*g*_s_) and the photochemical yield of photosystem II (Φ_PSII_) under saturating light conditions, with 400 µmol CO_2_ mol^-1^ air at 25 ºC. The electron transport rate (*ETR*) was calculated as in Carriquí et al (2015) using light curves conducted under low O_2_ conditions to estimate the alphabeta compound (Valentini *et al*., 1995). Light respiration (*R*_light_) was considered as half of the dark respiration rate, with plants having been exposed to darkness for at least 30 minutes (Niinements *et al*., 2005). From these measurements, mesophyll conductance to CO2 diffusion (*g*_m_) was estimated using the variable *J* method of Harley *et al*. (1992). For ferns, *C. japonica, A. thaliana* and *D. antarctica*, the CO_2_ compensation point in the absence of respiration (*Γ*\*) were assumed to be 101.93, 42.75, 50.22 and 34.83 μmol mol^-1^, respectively (Gago *et al*. 2013; Hanba *et al*., 1999; Whitney *et al*., 2011; Sáez *et al*., 2017).

### Anatomical characterization

After gas exchange measurements, small pieces of *D. antarctica* leaves enclosed in the leaf cuvette were fixed under vacuum pressure using a solution of 4% glutaraldehyde and 2% paraformaldehyde in a 0.01 M phosphate buffer (pH 7.4). The samples were then post-fixed for 2 hours in 2% buffered osmium tetroxide, dehydrated through a graded ethanol series, and embedded in LR white resin (London Resin Company). They were kept in an oven at 60 ºC for 48 h (Tosens *et al*., 2015).

Semi-fine and ultra-fine cross sections (0.8 μm and 90 nm, respectively) were cut using an ultramicrotome (Leica UC6, Vienna, Austria). Semi-fine sections were stained with 1% toluidine blue and observed with an Olympus BX60 light microscope. Images were captured at 200X magnification using a digital camera (U-TVO.5XC; Olympus, Tokyo, Japan). From these images, leaf thickness (*T*_leaf_) was measured. Ultra-fine cross sections were contrasted with uranyl acetate and lead citrate and photographed at 30000X magnifications using a transmission electron microscope (TEM H600; Hitachi, Tokyo, Japan). From these pictures, cell wall thickness (*T*_cw_) was calculated. Additionally, a cell curvature correction factor was determined using an average length/width ratio from five randomly selected cells (Thain, 1983). For each parameter, final values were obtained averaging 10 measurements from randomly selected cell structures using ImageJ software (Wayne Rasband/NIH, Bethesda, MD, USA).

### Cell wall composition analyses

Cell wall composition analyses were performed on the same leaves used for gas exchange measurements or in neighboring ones. Plants were kept in darkness overnight to minimize foliar starch accumulation. The next morning, approximately 700 mg of fresh leaf tissue were cut into small portions and were placed in screw-capped glass tubes filled with absolute ethanol. Samples were boiled until bleached, and cleaned with >95% acetone to obtain the alcohol-insoluble residue (AIR), which approximates the total cell wall amount. All AIRs were digested with α-amylase to remove starch residues. Once no starch residues were detected, three analytical replicates of each AIR weighing around 3 mg were hydrolysed with 2 M trifluoroacetic acid at 121 °C for 1 hour. They were then centrifuged at 13000 *g*, resulting in an upper aqueous phase (non-cellulosic cell wall fraction) and a pellet (cellulosic cell wall fraction). Supernatants were stored at -20 °C for hemicellulose and pectin quantifications. Pellets were washed with distilled water and >95% acetone, air-dried at room temperature, resuspended in 200 μl of 72% (w/v) sulfuric acid for 1 hour, diluted to 6 ml with distilled water, and heated at 121°C until degraded. After cooling, the resulting aqueous solutions were used for cellulose quantification. Cellulose and hemicellulose concentrations were determined using the phenol-sulfuric acid method (Dubois *et al*., 1956) from a glucose calibration curve. Pectin quantification followed the protocol by Blumenkrantz & Asboe-Hansen (1973) from a galacturonic acid calibration curve. All measurements were conducted using a Multiskan Sky Microplate Spectrophotometer (ThermoFisher Scientific).

### Statistical analysis

Data were processed in R (version 4.3.3) (R Core Team, 2023) using *ggplot2* and *smatr*. Standard Major Axis (SMA) regression was applied for linear relationships. For non-linear fittings, the function nls () from the “stats” package were used.

---

**Fig. S1.**
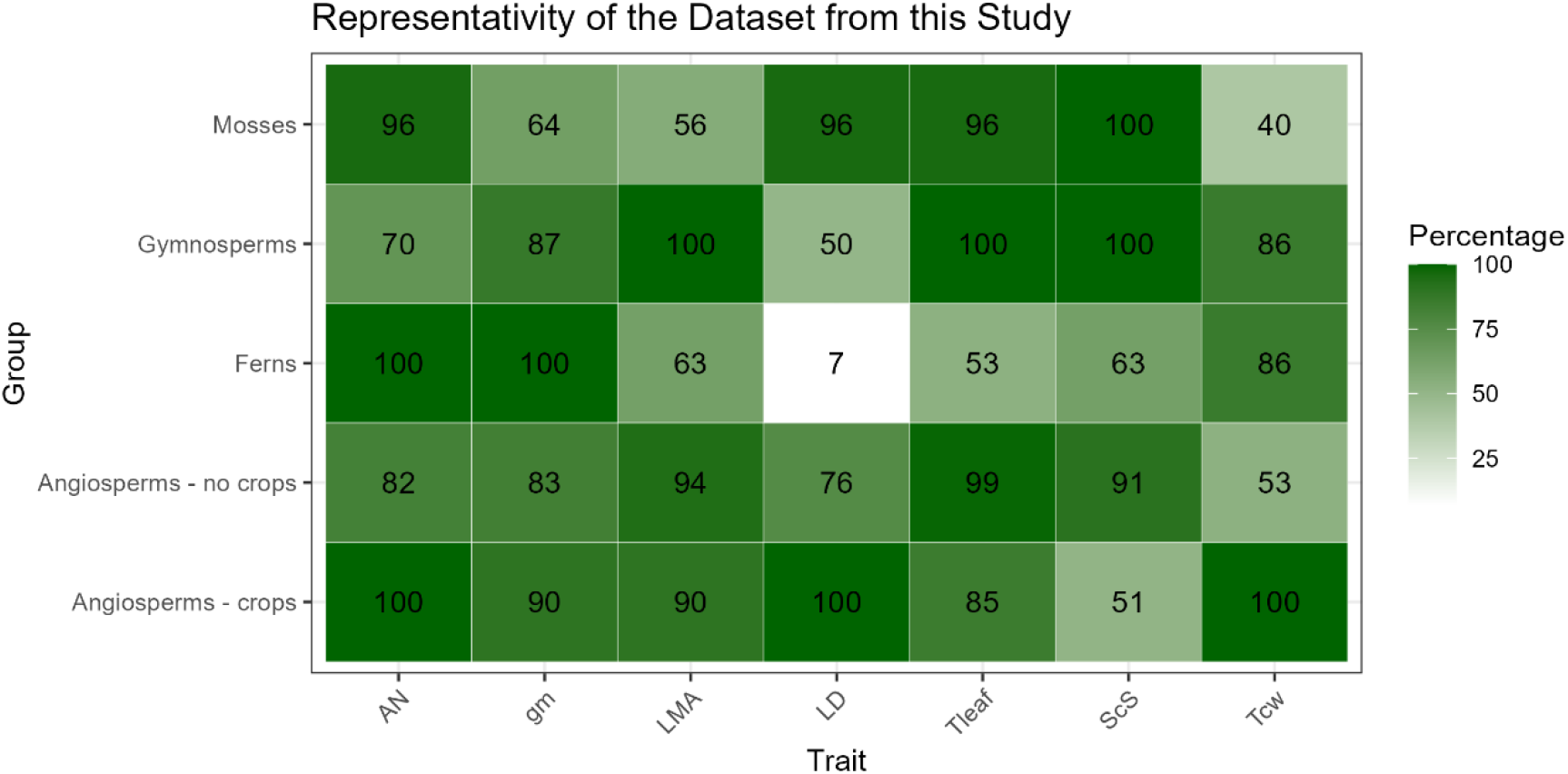
Heatmap showing the percentage of the published range (excluding the first and last decile) covered for each trait and phylogenetic group considered. Traits include A_N_ (net assimilation), g_m_ (mesophyll conductance), LMA (leaf mass per area), LD (leaf density), T_leaf_ (leaf thickness), S_c_ /S (chloroplast surface area exposed to intercellular airspaces per unit leaf area), and T_cw_ (cell wall thickness). The colour gradient from white to dark green represents increasing percentages, with 100% being the best coverage. Previously published range was considered from the data compilation from Huang et al. (2022) for gymnosperms, ferns and angiosperms—both crops and wild species—, while for mosses data from Carriquí et al. (2019), Perera-Castro and Flexas (2022), Roig-Oliver et al. (2021b), Bansal et al. (2012), and Wang et al. (2016) were considered.

**Fig. S2.**
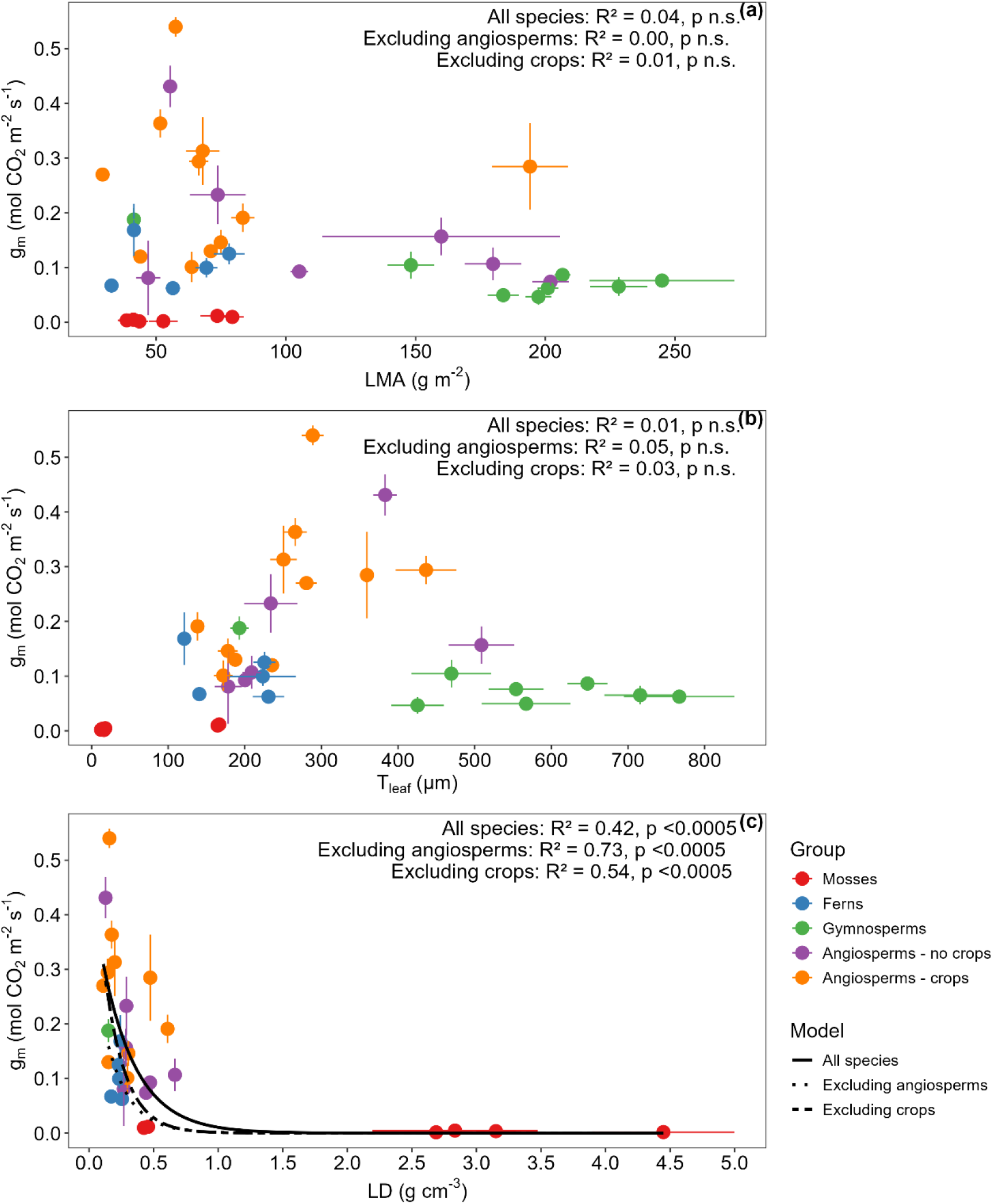
Relationships among mesophyll conductance to CO_2_ diffusion (g_m_) in relation to (a) leaf dry mass per unit area (LMA), (b) leaf thickness (T_leaf_) and (c) leaf density (LD) in C_3_ plants. Lines represent (a,b) linear and (c) exponential decay fittings on data pooled across all phylogenetic groups, excluding crops and excluding all angiosperms. Significant lines in (c) are exponential decay regression lines. Data was compiled from Carriquí et al. (2020), Clemente-Moreno et al. (2019), Nadal et al. (2020), Nadal et al. (2023), Roig-Oliver et al. (2020b,c;2021b,c;2022), and from newly-measured species.

**Fig. S3.**
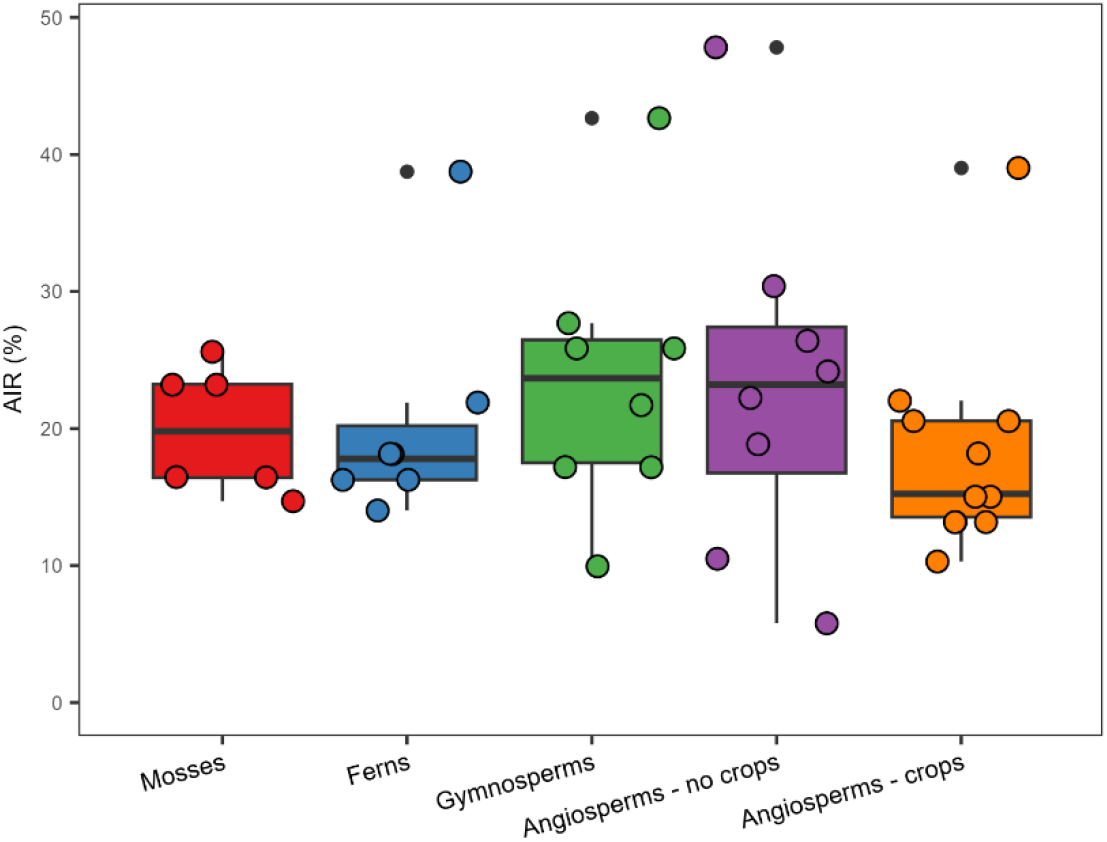
Dot boxplots showing the mass of cell walls prepared as alcohol insoluble residues per leaf dry weight (AIR) for mosses (n = 6), ferns (n = 8), gymnosperms (n = 8), wild angiosperms (n = 10), and angiosperm crops (n = 11). Data was compiled from Carriquí et al. (2020), Clemente-Moreno et al. (2019), Nadal et al. (2020), Nadal et al. (2023), Roig-Oliver et al. (2020c), Roig-Oliver et al. (2020b), Roig-Oliver et al. (2021c), Roig-Oliver et al. (2021b), Roig-Oliver et al. (2022), and from newly-measured species.

## Notes

### Competing Interest Statement

The authors have declared no competing interest.

